# Serial invasions can disrupt the time course of ecosystem recovery

**DOI:** 10.1101/2021.10.29.466526

**Authors:** Vadim A. Karatayev, Lars G. Rudstam, Alexander Y. Karatayev, Lyubov E. Burlakova, Boris V. Adamovich, Hanna A. Zhukava, Kristen T. Holeck, Amy Lee Hetherington, James R. Jackson, Christopher W. Hotaling, Tatyana V. Zhukova, Tamara M. Mikheyeva, Raisa Z. Kovalevskaya, Oleg A. Makarevich, Darya V. Kruk

## Abstract

The impacts of species invasions can subside or amplify over time as ecosystems “adapt” or additional invaders arrive. These long-term changes provide important insights into ecosystem dynamics. Yet studies of long-term dynamics are rare and often confound species impacts with coincident environmental change. We synthesize many-decade time-series across ecosystems to resolve shared changes in seven key features following invasion by quagga and zebra mussels, two widespread congeners that re-engineer and increasingly co-invade freshwaters. Six polymictic shallow lakes with long-term data sets reveal remarkably similar trends, with the strongest ecosystem impacts occurring within 5-10 years of zebra mussel invasion. Surprisingly, plankton communities then exhibited a partial, significant recovery. This recovery was absent, and impacts of initial invasion amplified, in lakes where quagga mussels outcompeted zebra mussels and more completely depleted phytoplankton. Thus, invasion impacts subside over time but can amplify with serial introductions of competing, even closely similar, taxa.

## Introduction

Introductions of keystone or ecosystem engineering species can profoundly transform multiple ecosystem features^1^, ^2^. For example, cattle can overgraze grasslands^3^, wolf re-introduction can allow grassland conversion into forests^4^, and zebra mussel invasions can transform lakes from turbid to clear water phases^1,5,6^. Many species introductions are intentional (e.g., for biological control), with consumer or predator re-introductions becoming a widespread ecosystem restoration strategy. Simultaneously, ongoing unintentional introductions of invasive species are progressing across regions and can cause socioeconomically undesirable ecosystem shifts. Reversing invasion impacts requires costly or risky management interventions (e.g., biological control^7^). Both management issues raise the question: how strong are the ultimate impacts of widespread species introductions and how quickly do they affect different ecosystem features?

The response of ecosystems to a species introduction can be complex and difficult to predict in a given system. A parsimonious null hypothesis is that a species’ impacts increase monotonically to a maximum that is maintained over time. Alternatively, initial impacts of introductions can subside as species traits or ecosystem structure adapt (reviewed ^8^); these changes could vary widely depending on environment or makeup of resident species^8–10^. Population abundance and impacts of invasive species often exhibit an invasion cycle where after initial high abundance, invader populations decline^11–13^ due to changes in community structure, density-dependent changes in the abundance of the invader, evolutionary and behavioral adaptations by resident species to the new invader (learning, eco-evo dynamics), or other factors.

For invasive species, the time course of ecosystem impact and recovery may break down if multiple species are introduced in series (e.g., invasion meltdown^14^). Interactions among invaders have been studied extensively for plants^15^, aquatic, and terrestrial animals^16,17^. These authors concluded that the ecological impact of an additional invader is hard to predict, and can be neutral, synergistic or antagonistic. In general, invasion biology anticipates small additional impacts of invaders when they are functionally similar to species already present in an ecosystem^8,18,19^. Alternatively, competition theory^20^ suggests greater impacts from a second invader that outcompetes an initial invader by utilizing ecosystem resources more efficiently and completely (*reviewed* ^21^).Therefore, even for functionally similar species the predicted impacts of serial invasions range from minimal ecosystem change to an amplification of initial invader’s impacts and prevention of ecosystem recovery.

Zebra mussels (*Dreissena polymorpha*, Pallas 1771) and quagga mussels (*D. rostriformis bugensis*, Andrusov 1897) exemplify widespread invaders that increasingly co-invade waterbodies. Both species have high reproductive and dispersal potential, often comprise a large portion of animal biomass in invaded ecosystems^12,22–24^, and represent perhaps the most aggressive freshwater invaders^12^. Whilst zebra mussels exhibit faster landscape-level spread, quagga mussels increasingly invade waterbodies that already had established zebra mussel populations^12,25^. As highly efficient suspension feeders, dreissenid species are both powerful ecosystem engineers and restructure energy flows from pelagic to benthic habitats^1,26–30^. Yet the magnitude, pace, and interaction of impacts caused by introduced species such as dreissenids remain obscure because long-term ecosystem studies are rare and analyzed separately, which confounds invasion impacts with any coincident environmental changes.

Here, we assemble high-resolution data sets spanning seven key ecosystem features, five decades, and six lakes in three regions (New York State, Belarus, and The Netherlands) invaded by dreissenid mussels. We analyze changes shared among lakes across time since invasion to robustly attribute ecosystem changes to the effects of dreissenid introduction rather than system-specific environmental changes. We begin with the question: how quickly do zebra mussel impacts manifest in each ecosystem feature? Next, we test the null expectation that invasion impacts increase monotonically to a maximum level *versus* the alternative hypothesis that impacts decline after an initial peak due to complex invasion dynamics or when native communities adapt to the invader. Then, in systems experiencing additional quagga mussel invasions, we evaluate whether serial invasions of competing species magnify the impacts of the initial invader.

## Results and Discussion

We study long-term trends collected in six polymictic lakes located in Belarus, the Netherlands, and New York, USA (Table 1). These shallow freshwater ecosystems have long time series of zebra mussel impacts (20-37 years) on both benthic and plankton communities, with an additional 6-15 years of baseline pre-invasion data in each system. These datasets are among the longest dreissenid invasion data series available worldwide^25^. We analyze changes in four communities (phytoplankton, zooplankton, zoobenthos, macrophytes) and three abiotic variables (Secchi depth, chlorophyll, phosphorous) as a function of time since the initial invasion by zebra mussels and the subsequent (serial) invasion by quagga mussels. All lakes have a strong long-term dreissenid presence, but span a wide range of mean mussel biomass (100-800 g/m^2^), lake morphometry (1.8-9 m mean depth), and area (15-200 km^2^).

**Table 1.** Physical characteristics of Lake Lukomskoe^70^, the Narochanskie Lakes^71^, Oneida Lake^46^, and lakes Veluwe and Eem^52^ with the dates of *Dreissena* spp. first records. Mussel filtration capacity denotes percent of lake volume filtered per day at 20°C (40ml hr^−1^ g^−1^, ^69^ filtration in Veluwe and Eem from Noordhuis et al.^52^ and year(s) corresponding to the estimate.

| Lake | Coordinates | Surface area, km<sup>2</sup> | Maximum depth, m | Average depth, m | *D. polymorpha* introduction | *D. r. bugensis* introduction | Zebra mussel biomass (g/m<sup>2</sup>); filtration capacity | Quagga mussel biomass (g/m<sup>2</sup>); filtration capacity |
| --- | --- | --- | --- | --- | --- | --- | --- | --- |
| Lukomskoe | 54°40'N, 29°5'E | 36.7 | 11.5 | 6.7 | 1972 | None | 262; 3% (1970-1990) |  |
| Naroch | 54°51'N, 26°45'E | 79.6 | 24.8 | 9.0 | 1989 | None | 114; 1.5% (1993-2016) |  |
| Myastro | 54°51'N, 26°53'E | 13.1 | 11.3 | 5.4 | 1984 | None | 409; 7% (1993-2017) |  |
| Oneida | 43°10'N, 75°52'W | 207.0 | 16.8 | 6.8 | 1991 | 2005 | 804; 12% (1992-2006) | 1178; 15% (2007-2018) |
| Veluwe | 52°24'N, 5°45'E | 31.3 | 5.0 | 1.8 | 1995 | 2009 | NA; 40% (2000-2010) | NA; 77% (2013) |
| Eem | 52°17'N, 5°20'E | 15.2 | 4.0 | 2.1 | 1995 | 2009 | NA; 66% (2000-2008) | NA; 429% (2011, 2013) |

### Lake ecosystem impacts manifest quickly

Across these six ecosystems and every feature examined, we find strong ecosystem changes that peak within 5 to 10 years of the initial zebra mussel invasion (Fig. 1). In lakes with the longest time series of dreissenid population dynamics, zebra mussel biomass increased to a steady level within 3 years (Naroch, Oneida), although a partial decline followed this initial increase in Lake Lukomskoe (Fig. 2). Our best-fit splines of each ecosystem variable *versus* time since invasion fitted to cumulative data from all lakes aptly summarize trends within each lake (mean consistency C=0.7; *see* Methods). These trends identify significant declines in total phosphorus, chlorophyll *a*, phytoplankton biomass, and zooplankton biomass accompanied by increases in Secchi depth, zoobenthos, and macrophyte coverage.

**Figure 1.**
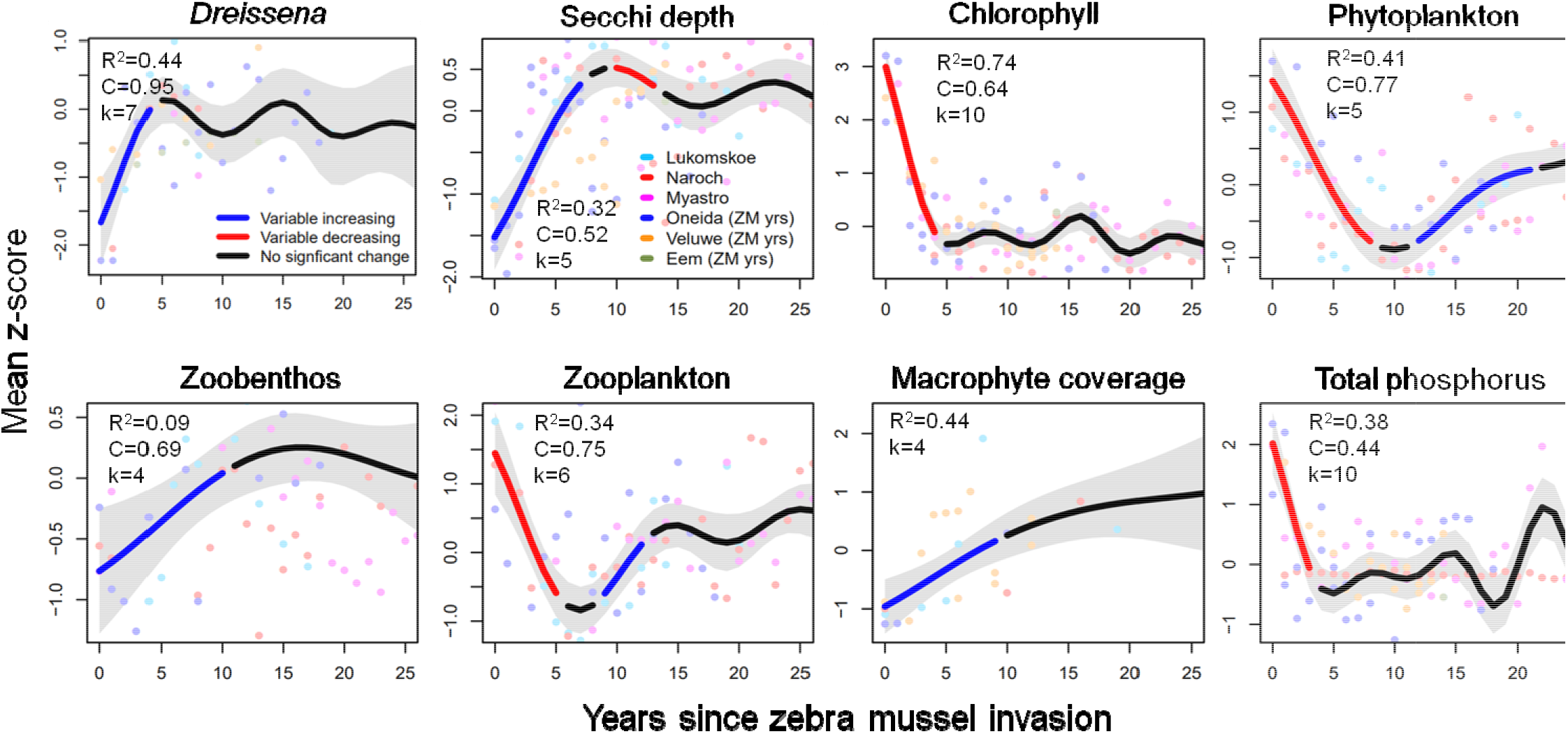
Zebra mussel invasion corresponds to significant long-term changes in ecosystem variables across six polymictic lakes in Europe and North America. Lines denote mean z-score changes in variables predicted by best-fit splines across years since invasion (x-axes) across all lakes under conditions of 100% zebra mussel dominance. For each variable, line colors denote significant increases in blue, significant decreases in red, no significant change in black, and gray areas denote 95% confidence intervals of the mean. For each variable, R^2^ denotes the proportion of total variance explained by fitted models, *k*_*x*_ denotes the basis dimension (maximum smoothness) of fitted splines, and C denotes the consistency of lake-specific trends (Fig. S1) with the cumulative trend plotted (see *Methods*). Note, only 20 years of post-invasion data from Lake Lukomskoe was used in model fitting as subsequent hypoxia depleted zebra mussel biomass. Observations (points) under conditions of quagga dominance omitted as they did not inform model projections shown.

**Figure 2.**
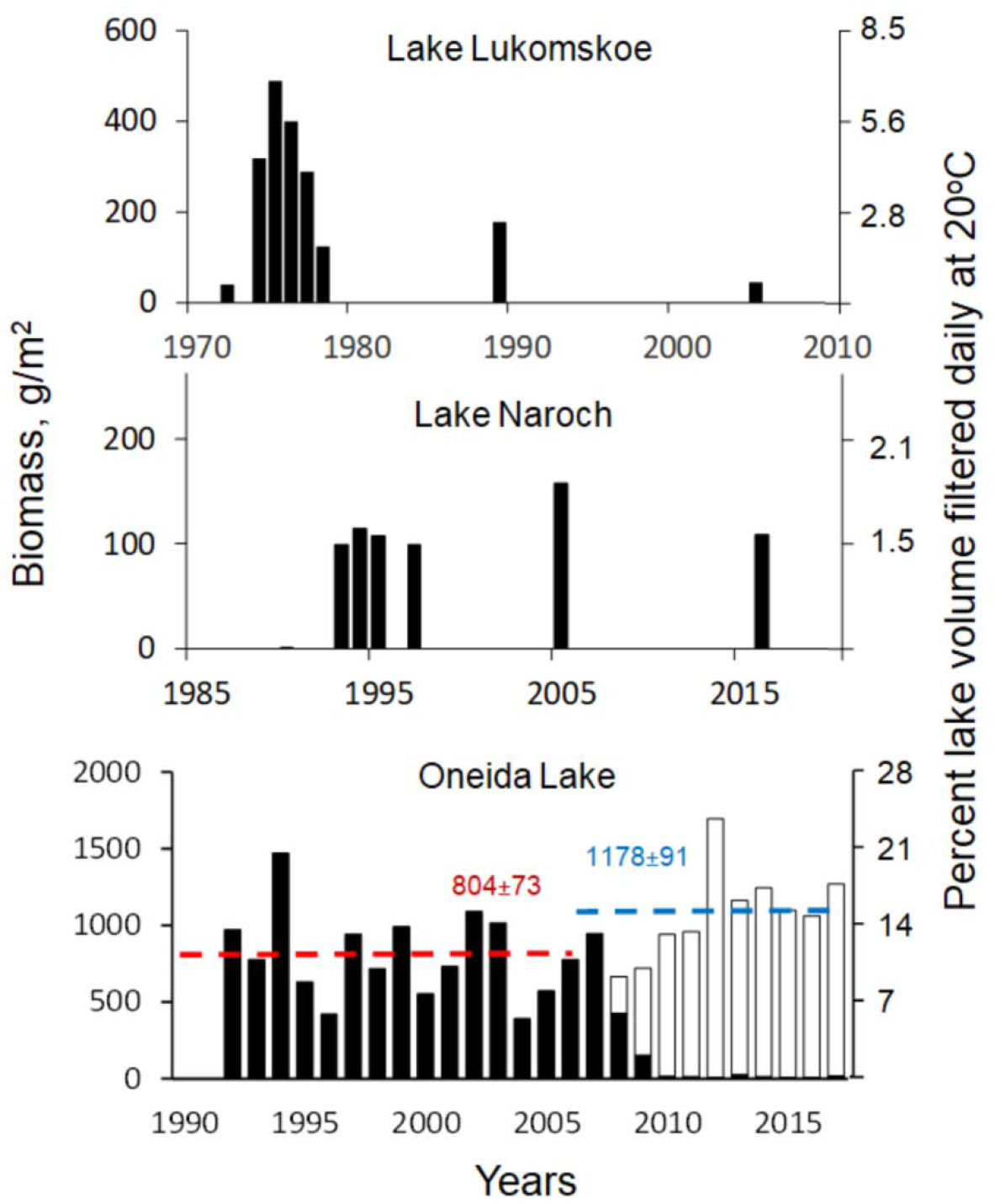
Long-term dynamics of wet biomass of zebra mussels (black bars) and quagga mussels (white bars) in lakes Lukomskoe, Naroch, and Oneida Lake. Horizontal dashed lines for Oneida Lake denote mean mussel biomass +/− standard error for periods of zebra mussel dominance (1992 – 2006, red line) and quagga mussel dominance (2009 – 2017, blue line). Secondary axes denote the percent of lake volume filtered daily, estimated from biomass, mean depth, and 40ml hr^−1^ g^−1^ filtration rate^69^.

These changes characterize a shift from a turbid to a clear water phase (“benthification” ^5^) and a net transfer of resources from the plankton community to the benthos by mussel filter feeding (reviewed^1,27,28,30,31^). Mussels directly reduced chlorophyll (R^2^=0.74, C=0.64) and phytoplankton (R^2^=0.41, C=0.77), in turn reducing zooplankton (R^2^=0.34, C=0.75). Mussels have a positive but less consistent, indirect impact on zoobenthos and facilitate macrophyte coverage by deepening the photic zone. In one lake we did not detect a decline or recovery in zooplankton after zebra mussel invasion (Oneida Lake, see Fig. S1 for lake-specific trends). Zooplankton dynamics in this period were highly variable due to multiple factors, including fish predation; in Oneida Lake age-0 fish are known to affect zooplankton biomass ^32^ and relatively low age-0 yellow perch abundance occurred after the zebra mussel invasion ^33^. Lower zebra mussel impacts on zooplankton in Oneida Lake may also have been influenced by the deepened photic zone, which in this lake compensated for reduced phytoplankton biomass, producing little change in total phytoplankton primary production^34^.

Rooted in high-resolution, ecosystem-level monitoring studies, these results support findings across aquatic and terrestrial systems. Bradley et al.^35^ found that invaders have disproportionally strong impacts on species they consume (here, phytoplankton) at low levels of invader abundance. In line with this, Strayer et al.^8^ highlight the importance of considering both the short-term, ‘acute impacts of invaders as well as long-term, ‘chronic’ invasion outcomes.

### Ecosystems partially recover from invasion

Following initial invasion impacts, plankton communities did not stabilize and instead partially recovered in ecosystems with only zebra mussels (Fig. 1). Three lakes exhibited significant, synchronized recoveries towards pre-invasion levels of phytoplankton biomass (~50%, C=0.77), zooplankton biomass (~30%, C=0.75), and Secchi depth (~15%); Secchi depth recovery was weaker because this feature was less synchronized across lakes (C=0.52; see *Results robustness* below).

Two broad possible drivers can underlie these partial plankton community recoveries: (a) reduced grazing capacity of the mussel population and (b) increased phytoplankton resistance to grazing. A decline in mussel biomass or filtering activity could arise through intraspecific processes, for example as competition reduces mussel populations^36^ or as populations become dominated by large adults with lower mass-specific filtration. Mussel biomass could also decline as species interactions or stochastic events increase mortality. These include anoxia^37^, native predators learning to feed on mussels^38,39^, or the arrival of new predators^40^. However, the declines in mussel impacts we observe happened synchronously across lakes and are not consistent with stochastic events such as new invaders or anoxic conditions. Demographic processes within the invading population and functional or numerical responses by native molluscivores may operate on similar time scales across systems, and are therefore a more likely driver of observed recoveries.

An alternative mechanism for the partial plankton community recoveries is increased resistance of phytoplankton to grazing. This can arise with a change in traits or community structure. For example, phytoplankton communities can change to species that avoid bottom filter feeders by regulating their buoyancy or are too large to be consumed by mussels, such as some cyanobacteria^41,42^. New studies and data are needed to test this hypothesis, which may additionally explain why phytoplankton biomass increased while chlorophyll *a* concentration remained stable in each of the three lakes not invaded by quagga mussels (Fig. 1, S1). This result implies that the phytoplankton community changed towards species with less chlorophyll per unit biomass, perhaps an adaptation to higher light levels^43^ ^44^.

In other systems, it is known that ecosystem structure and community composition can shift towards a higher abundance of animals consuming the introduced species or shift towards taxa resistant to the introduced species’ impacts. Therefore, the impacts of introduced species commonly change over time due to evolution, shifts in species composition, or abiotic changes in the environment (reviewed^2,8^). For example, invasive garlic mustard evolved reduced allelochemical concentration, while several native species evolved increased tolerance of garlic mustard allelochemical (reviewed^2^). Introduction of domestic ungulates leads to a dominance of grazing- and trampling-resistant grasses or woody vegetation^45^). In most instances, such changes in species impacts can only be resolved by long-term studies^2,8^). The growing emphasis on long-term monitoring could provide a temporal context for species introductions and insights into the common mechanisms by which ecosystems respond to invasions.

### Serial invasions amplify ecosystem impacts

Approximately 14 years after the initial zebra mussel invasion, lakes Oneida, Eem, and Veluwe experienced a second invasion by quagga mussels, which quickly displaced zebra mussels to low densities (5 years in Oneida^46^). We find that this serial invasion increased total mussel biomass and amplified the ecosystem impacts of the preceding zebra mussels (Fig. 3). Specifically, quagga mussels further depleted chlorophyll *a* concentrations and prevented the recovery of phytoplankton biomass observed in lakes experiencing a single invasion. In Oneida Lake, quagga mussel invasion also strongly reduced zooplankton biomass, an effect we did not detect following the initial zebra mussel invasion.

**Figure 3.**
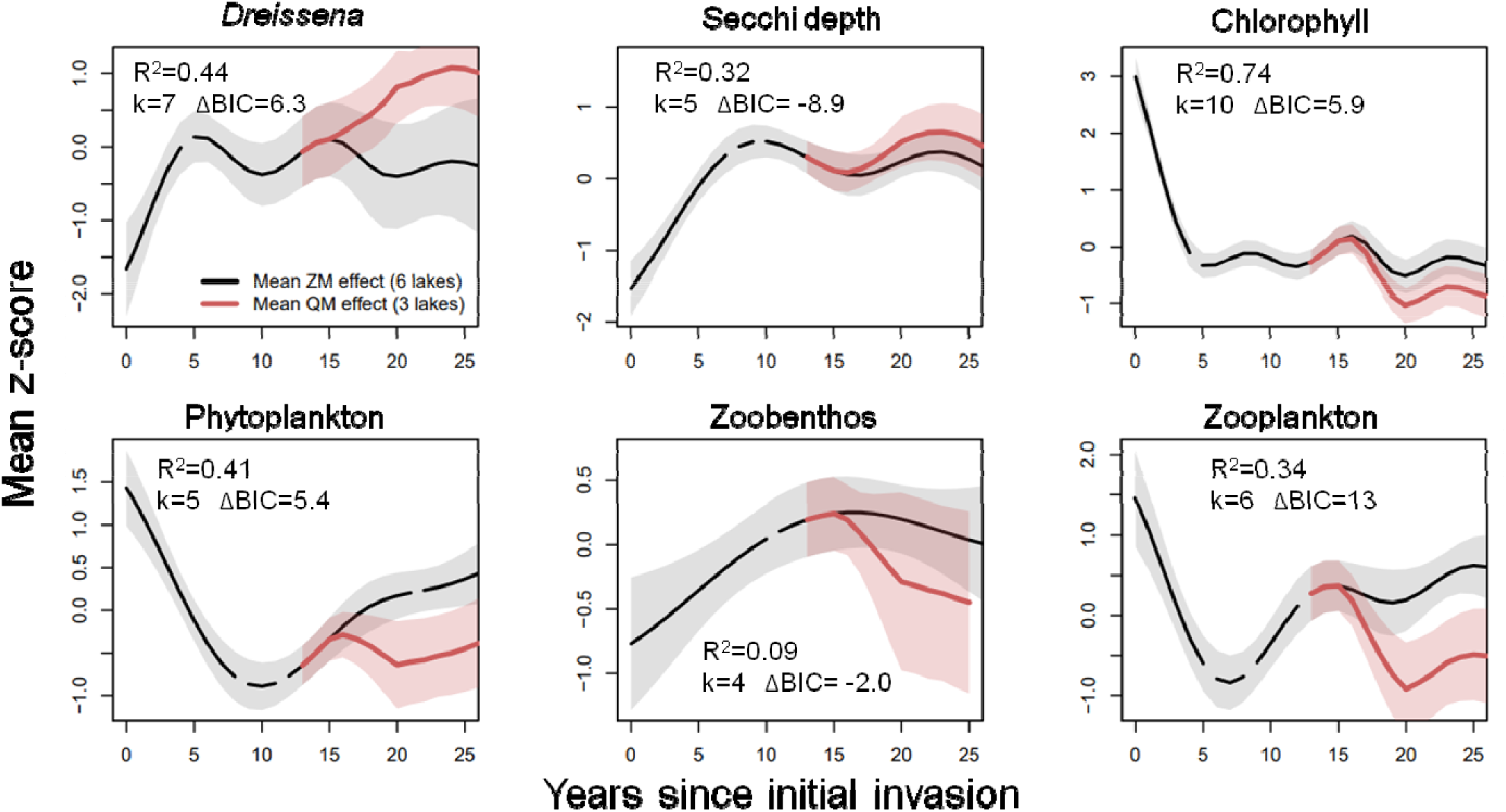
Secondary invasion by quagga mussels leads to significant changes in several ecosystem variables across three polymictic lakes. Lines denote mean z-score changes in variables predicted by best-fit splines with only zebra mussels (black lines as in Fig. 2) and with the added invasion of quagga mussels at year 15 (red lines, lakes Eem, Veluwe, and Oneida). Following data in Oneida Lake, the proportion of zebra mussels in models with quagga mussels was decreased to ~0% by year 20. Shaded areas denote 95% confidence intervals of each mean. R^2^ denotes the proportion of total variance explained by fitted models and k denotes the maximum basis dimension (decreasing smoothness) of fitted splines. Note that data on zoobenthos, phytoplankton, and zooplankton with quagga mussels are available only for Oneida Lake. Averages are from May through October.

Serial *Dreissena* invasions amplify ecosystem impacts because quagga mussels utilize resources more completely than zebra mussels across space and time. Quagga mussels utilize a wider array of habitat types because they colonize rocky shallows as well as colder, soft-sediment habitats in deep areas of lakes (reviewed^12,36^). This allows the species to achieve higher total biomass, as evident in the North American Laurentian Great Lakes where serial quagga mussel invasion increased total dreissenid biomass ten-fold^36^). In addition to higher biomass, in Oneida Lake we find that quagga mussels also utilize resources more completely across seasons by actively filter-feeding during cold periods. Compared to years of zebra mussel dominance (1993-2007), quagga mussel dominance in 2009-2017 corresponded to significantly greater visibility and lower chlorophyll concentrations during early-spring and late-fall but not in warmer months (Fig. 4b, c). This helps explain findings in lakes Michigan and Huron, where the spring diatom blooms disappeared only after quagga mussel invasion^12,47–49^. These patterns likely arise from higher physiological activity of quagga mussels compared to zebra mussels at colder temperatures^50,51^ and explain the higher per-biomass impact of the species on phytoplankton (Fig. 4a).

**Figure 4.**
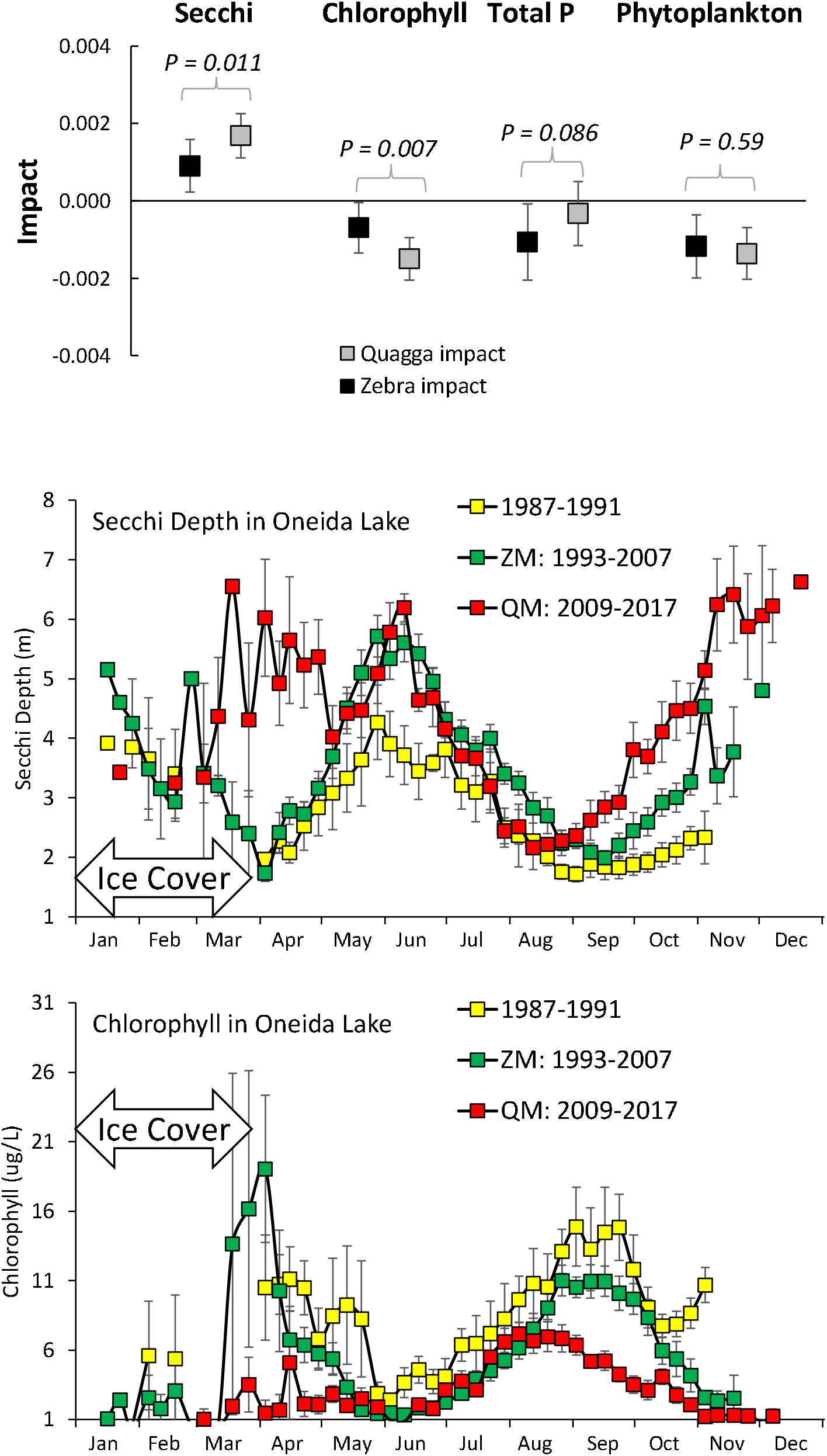
Compared to zebra mussels, quagga mussels in Oneida Lake have greater per-biomass impacts on visibility and chlorophyll (a) due to filter feeding during cold months (b, c). In points show the effect sizes of each species in general linear regression models of Secchi depth, chlorophyll *a*, total phosphorous and phytoplankton across 1987-2017 (averages from March through November) as a function of zebra mussel biomass (black) plus quagga mussel biomass (gray). *P*-values for each ecosystem feature denote the presence of significant differences in coefficients between the two species in linear hypothesis tests, and error bars denote +/− 2 standard errors of each coefficient. (b and c) show weekly dynamics of chlorophyll *a* averaged (± 1 standard error) across years before *Dreissena* invasion (1987-1991), during zebra mussel dominance period (1993-2007), and during quagga mussel dominance period (2009-2017).

Taken together, our results suggest that in displacing zebra mussels, quagga mussels achieve higher total biomass (Fig. 2; see Nordhuis et al. ^52^ for the two Dutch lakes) as well as higher per unit biomass impacts (Fig. 4), and therefore deplete phytoplankton more completely. Classical resource competition theory predicts this to be a general phenomenon when the serially introduced species are competitors: a species establishes and outcompetes a preceding invader because it can deplete resources more completely (i.e., persist at a lower R*,^20^). Thus, competitive displacement by functionally similar invasive species (often from the same genus) is known as “over-invasion” and has been documented for wasps^53^, mosquitoes^54^, foxes^55^, nectar-thieving ants^56^, and amphipods^57^; reviewed^21^. Our results demonstrate that even for closely related species, subtle differences in environmental preferences (habitat, temperature) drive large ecosystem-level impacts of serial invasions.

### Results robustness

Initial and serial invasion impacts found here are remarkably consistent across the shallow, polymictic lakes studied here even though these ecosystems span an 8-fold range in long-term *Dreissena* biomass, 14-fold range in lake area, and 4-fold range in mean lake depth. The different regions and dates of initial invasion (1972-1995) also mean that any undetected changes in the abiotic environment could only skew our qualitative results if they happened at similar times since invasion across all lakes, which is unlikely. However, we caution against using our results as a quantitative roadmap of dreissenid invasion impacts for two reasons.

First, trajectories in any single system may deviate from aggregate lake trends in our analyses due to system-specific trajectories in mussel biomass and the abiotic environment mediating mussel impacts. The negative impacts of mussel filter feeding on phytoplankton might be offset by increased nutrient loading into a lake (e.g., urbanization, increased fertilizer use) or, potentially, exacerbated by concurrent efforts to reduce nutrient loading. Excessive organic pollution that depletes dissolved oxygen is also known to reduce or even extirpate *Dreissena* populations, including in the central basin of Lake Erie^37^ and in Dutch lakes during the 1970s^6,52^. Indeed, the 1995 ‘re-invasion’ of lakes Veluwe and Eem by zebra mussels studied here occurred after large reductions in nutrient loading. Thus, the initial P declines in these lakes may be the results of both P loading reductions and the impact of mussels. Analogously, 10-fold decline in zebra mussel biomass observed in Lake Lukomskoe in 2005, 30 years after the peak in biomass (Fig. 2), was most likely driven by the increase in nutrient load and oxygen depletion caused by the fish hatchery launched in the lake in 1989^58^. This dramatic decline in zebra mussel population was associated with almost complete return of Lake Lukomskoe ecosystem to pre-invasion conditions (Fig. S2). Note that we only used the first 20 years of data for this lake in our analyses to avoid confounding our results with the increase in nutrient loading. Finally, quagga mussel biomass can exhibit very slow growth and reduced grazing in the profundal zones of deep lakes^36^, potentially limiting our quantitative insights to the shallow zones of the deep lakes.

Second, our approach may quantitatively under-estimate the degree of invasion impacts and ecosystem recovery. As systems differ in the precise timing of invasion impacts (increases, recoveries), the among-lake aggregate invasion impacts estimated here become dampened through an averaging effect. In the case of Secchi depth, for example, transparency peaked 5 years after invasion and then began to decline in lakes Veluwe and Oneida, but in Belarussian lakes Secchi depth continued increasing for 7-10 years after invasion (again, followed by a subsequent decline; Fig. S1). We expect differences in the timing of impacts to be the norm as, for example, the pace of *Dreissena* invasion slows in deeper, larger lakes^36^.

## Conclusions

Our findings show that the effects of species introductions on many ecosystem features can manifest quickly and partially subside, except when multiple species are serially introduced into a system. This insight comes from our decades-long perspective of ecosystem changes, which contrasts the short time span characteristic of most studies of invasive species that may only detect the increasing or decreasing trends depending on time since arrival. For other taxa, our approach shows how synthesizing monitoring studies can resolve introduced species’ impacts over time by focusing on shared changes across ecosystems *versus* time since introduction.

For managers, our results detail a comprehensive ecosystem-scale insight of *Dreissena* invasion impacts by first zebra and then quagga mussels. The changes in multiple ecosystem features resolved here also arise in studies of single systems^6,30,52,59^. As quagga mussels swiftly spread and colonize lakes already invaded by zebra mussels across Europe and North America (reviewed^12,60^), both initial and serial invasions confront managers with declines in phytoplankton, zooplankton, and potential declines in fisheries. In lakes Michigan and Huron, for instance, loss of spring diatom booms following serial *Dreissena* invasion was implicated in the decline of a burrowing amphipod (*Diporeia* spp.), and subsequent declines in commercially valuable lake whitefish (*Coregonus clupeaformis*) that feed on these amphipods ^47,48,61^. In these systems, declines in phytoplankton also correspond to declining salmon and lake trout populations^49^.

Our finding that serial invasion by a congener amplifies invasive species impacts shows that invaders can strongly affect ecosystems dominated by functionally similar species. This contrasts predictions of reduced ecological impacts of new invaders with low functional distinctiveness from existing species that emerge in meta-analyses^18,19^. Thus, we caution that functional or taxonomic distinctiveness can be a poor predictor of invasion impacts in a given system. This also re-emphasizes preventing invasive species’ spread as the most reliable means to minimizing their landscape-level impacts.

## Material and methods

### Study systems

For all Belarusian lakes and Oneida Lake we collected authors’ unpublished data as well as available information from peer-reviewed papers, unpublished reports and archive materials on water transparency (Secchi depth, m), total phosphorus concentration (mg/L), chlorophyll *a* concentration (mg/L), macrophyte coverage (%), wet phytoplankton biomass (g/m^3^), wet zooplankton biomass (g/m^3^), wet benthic biomass excluding molluscs (g/m^2^), and *Dreissena* spp. wet biomass (tissue and shells, g/m^2^; for detailed methods see Appendix S1). Note for Oneida Lake zoobenthos biomass was not recorded, and we used density instead. Information on the time course of chlorophyll, water clarity, macrophytes, and total phosphorus from lakes Eem and Veluwe was retrieved from Noordhuis et al.^52^.

### Zebra mussel ecosystem impacts

Throughout, we used a Generalized Additive Model (GAM) framework to resolve the impact of each species introduction on each ecosystem variable *x*. We first normalized observations in each lake *i* and year *t* into z-scores as 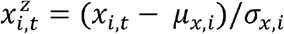. To account for both direct effects of invader biomass/activity and delayed ecosystem responses, we then model invasion effects on normalized variable time series *x*^*z*^ across all lakes as a smooth function of time using mgcv^62^ in R 3.5.2^63^. Specifically, to account for multiple potential invasions, we fit 2-dimensional splines to (1) years since zebra mussel invasion and (2) the relative abundance of zebra *versus* quagga mussel and assumed (and verified in preliminary analyses) a normal error distribution. While quagga mussels quickly became dominant in all lakes they invade ^25,64^, detailed mussel data were available only in Oneida Lake. However, the time dynamics of quagga mussels replacing zebra mussels is similar across lakes^64,65^ and we therefore assumed that relative dreissenid species abundance followed a similar pattern in the two Dutch lakes. More intermittent data on zebra and quagga mussel densities in the two Dutch lakes that suggest similar fast replacement of zebra mussels by quagga mussels^52^.

Given our focus on resolving general invasion impacts, we minimize lake-specific effects for each ecosystem variable *x* by using a Bayesian Information Criterion to select the maximum basis dimension *k*_*x*_ of the fitted spline (rejecting increases in *k*_*x*_ with ΔBIC<2). For each variable *x* we additionally fit separate lake-by-lake splines using the basis dimension *k*_*x*_ (omitting lakes with < 6 observations of *x*). We then calculate the total residual sum of squares across lake-specific splines tRSS_lake_, the residual sum of squares in the cumulative spline fitted to data from all lakes RSS_cuml_, and C = tRSS_lake_ / RSS_cuml_, an estimate of how well the cumulative spline captures lake-specific trends.

We approximated the timing and direction of zebra mussel effects based on the slope of fitted splines calculated using the package ‘tsgam’, where negative (positive) slopes indicate a decrease (increase) in the ecosystem variable. We conservatively determined changes as significant when (1) the confidence intervals of the slope at year *t* exclude zero and (2) the same change detection (i.e., positive or negative slope) is maintained for at least 4 consecutive years since zebra mussel invasion. For year “zero” (before zebra mussel invasion) we averaged data for Lake Lukomskoe for 1960-1970, Naroch 1978-1989, Myastro 1978-1987, Oneida Lake 1985-1989 and for lakes Veluwe and Eem 1985-1994.

### Effects of serial Dreissena invasions

Due to the limited time span of quagga mussel invasion, we resolved the impacts of this second invasion on each ecosystem variable by comparing the performance of our full models (i.e., 2d splines across time and invader species composition) and a null model of time only (i.e., 1d spline), with quagga mussel effects being significant for ΔBIC>4. We visualized quagga mussel effects by comparing the mean and 95% confidence intervals of each effect for (a) a lake invaded by zebra mussels only *versus* (b) a lake invaded first by zebra mussels at *t*=1 and then by quagga mussels at *t*=15 (by chance, the time of quagga mussel arrival since the zebra mussel (re)invasion was the same in lakes Eem, Veluwe, and Oneida).

To compare zebra and quagga mussel effects quantitatively, we measured the effects of each species’ biomass on ecosystem features in Oneida Lake, where biomass data are available annually for both species^46^. Unlike zebra mussels, substantial quagga mussel filtering activity and growth occurs in early spring and late fall^50^ ^66^. In this quantitative comparison only, we therefore included data from March – November and limited analyses to ecosystem features sampled over this entire period (Secchi depth, chlorophyll, total phosphorous, and phytoplankton), and included observations during 1987-1989 when both dreissenid species were absent from the lake. To measure impacts we used linear regressions of each ecosystem variable as a function of zebra mussel biomass plus quagga mussel biomass: 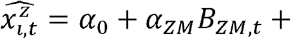 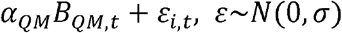. We then (1) evaluated the significance of each species’ impact (i.e., regression slope) and (2) compared the similarity of species’ impacts using linear hypothesis tests in the package ‘car’. Finally, to better visualize how differences in species effects arise, we compared the seasonal pattern of chlorophyll and Secchi depth in Oneida Lake in years with zebra mussel dominant and years with quagga mussel dominant.

## Acknowledgements

Oneida Lake surveys were supported by Cornell University with support from New York State Department of Environmental Conservation. We are grateful to Edward Mills for initiating the mussel studies on this lake and the many Cornell technicians and students supporting sampling over decades. Narochanskie Lakes surveys were supported by Belarusian State University and partially by the Belarusian Republican Foundation for Fundamental Research. The authors thank the researchers and technicians of Belarusian State University Naroch Biological Station, and graduate student I. Selivonchik who helped to collect and produce the data presented here, as well as divers V. Katulko and A. Soldatenkov for help in *Dreissena* sampling in Narochanskie lakes in 2016-2018. Visit to Cornell BFS for Adamovich and Zhukava was funded by Cornell University Biological Field Station.

## Author contributions

AYK, LGR, LEB, VAK, BVA, and HAZ conceived the study; VAK and LGR conducted analyses; VAK, LGR, AYK, and LEB wrote the paper, with input from other authors; all authors contributed data

## Materials & Correspondence

Lars Rudstam or Vadim Karatayev

## Supplementary Information

Additional supporting information may be found online at: [link to be added in production]

## Data Availability

Raw data are available from the Dryad Digital Repository: https://doi.org/10.5061/dryad.mpg4f4qzw^67^. Data from Oneida Lake also available at Knowledge Network for Biocomplexity: https://knb.ecoinformatics.org^68^.

