## Appendix S3: Fig. S2 for "Serial invasions can disrupt the time course of ecosystem recovery"

Karatayev, A. Y., Rudstam, L. G., Karatayev, V, A., Burlakova, L. E., Adamovich, B. V. Zhukava, H. A., Holeck, K. T., Hetherington, A. L., Jackson, J. R., Hotaling, C.W., Zhukova, T. V., Mikheyeva, T. M., Kovalevskaya, R. Z., Makarevich, O. A., Kruk, D. V. Serial invasions disrupt the time course of ecosystem recovery. Nature Communications.

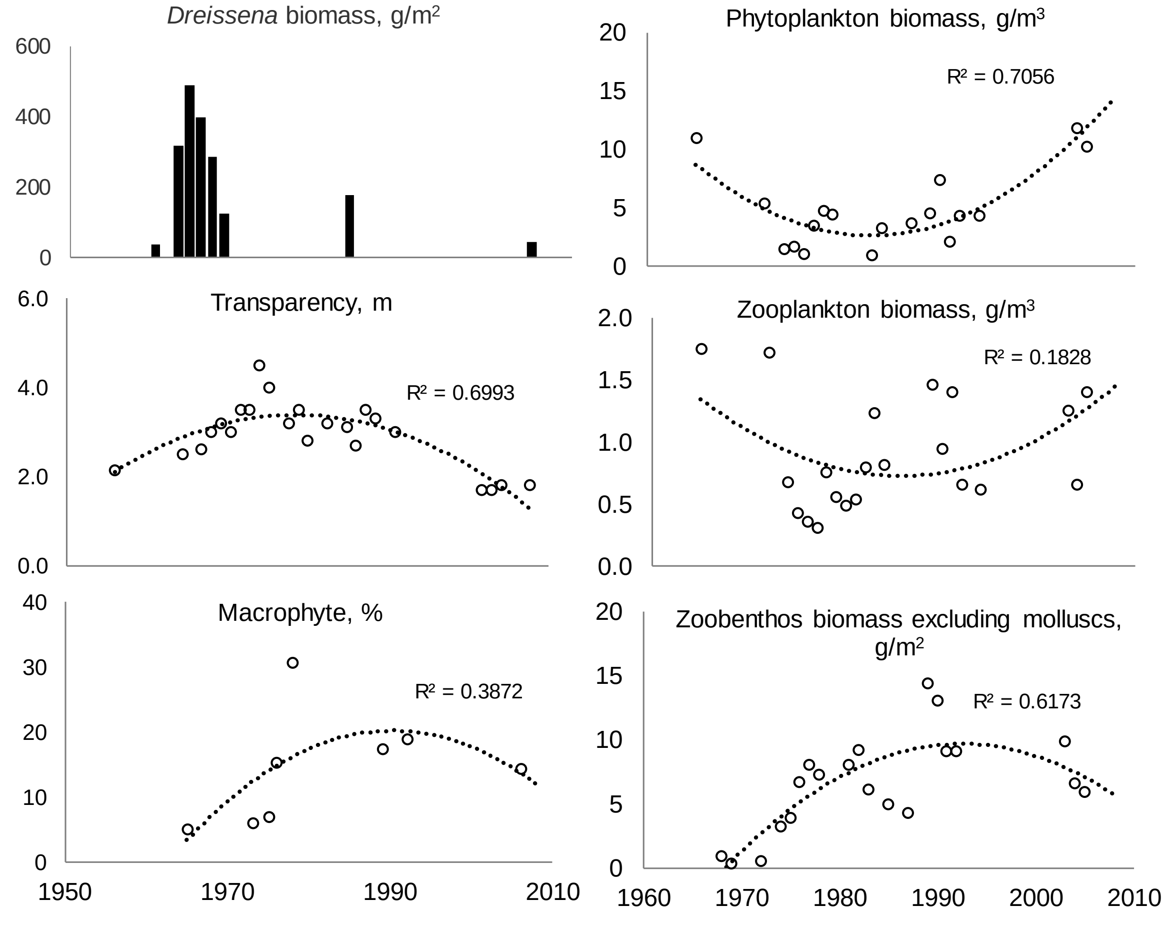


Figure S2. Long-term dynamics of *Dreissena polymorpha* and environmental variables in Lake Lukomskoe.
