## Appendix S2: Fig. S1 for "Serial invasions can disrupt the time course of ecosystem recovery"

Karatayev, A. Y., Rudstam, L. G., Karatayev, V, A., Burlakova, L. E., Adamovich, B. V. Zhukava, H. A., Holeck, K. T., Hetherington, A. L., Jackson, J. R., Hotaling, C.W., Zhukova, T. V., Mikheyeva, T. M., Kovalevskaya, R. Z., Makarevich, O. A., Kruk, D. V. Serial invasions disrupt the time course of ecosystem recovery. Nature Communications.

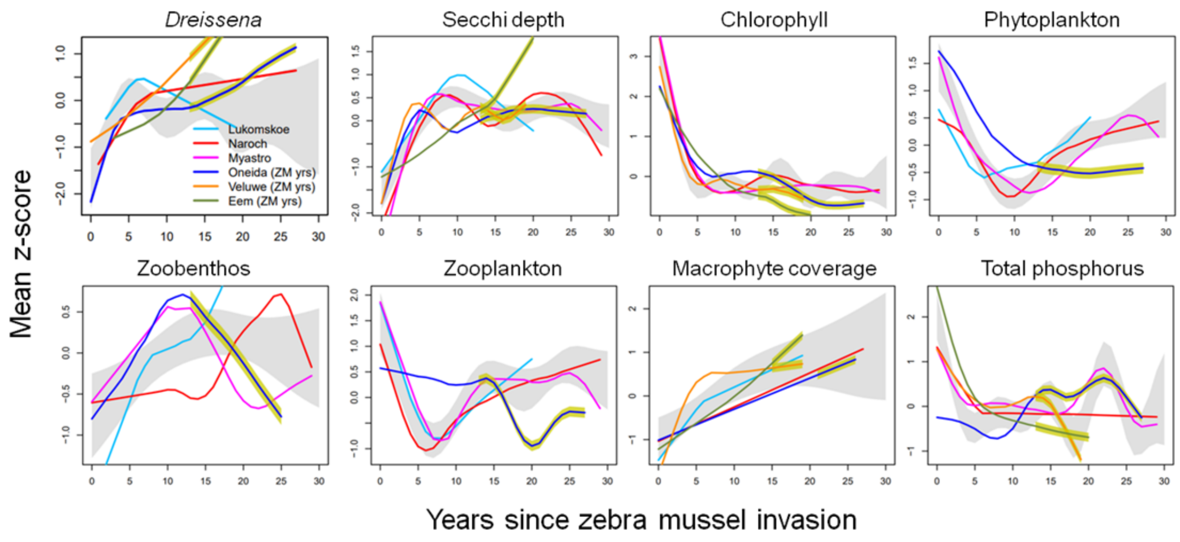


Figure S1. Smoothed lake-specific changes in Secchi depth (m), chlorophyll (µg/L), biomass of phytoplankton (g/m$^3$), zoobenthos (g/m$^2$), zooplankton (g/m$^3$), macrophyte coverage (%), and total phosphorus (µg/L), in polymictic lakes in Europe and North America following zebra mussel invasion. Gray areas indicate 95% confidence intervals of the mean zebra mussel effect. Red outlines around lakes Oneida, Eem, and Veluwe after year 15 denote ecosystem changes following quagga mussel invasions. Averages are from May through October.
