## Appendix S1 for "Serial invasions can disrupt the time course of ecosystem recovery"

Karatayev, A. Y., Rudstam, L. G., Karatayev, V, A., Burlakova, L. E., Adamovich, B. V. Zhukava, H. A., Holeck, K. T., Hetherington, A. L., Jackson, J. R., Hotaling, C.W., Zhukova, T. V., Mikheyeva, T. M., Kovalevskaya, R. Z., Makarevich, O. A., Kruk, D. V. Serial invasions disrupt the time course of ecosystem recovery. Nature Communications.

### METHODS

#### Study area and history of invasion

Studied lakes differed in the duration of *Dreissena* spp. invasion and the amount of information available. Lake Lukomskoe has the longest dataset of zebra mussel invasion (1972-2008) but has the least amount of available data, especially for the pre-invasion period. Due to the lack of primary data for Lake Lukomskoe we used published data, mostly summer or growing season averages (see below). Much more information (including primary data) is available for Narochanskie lakes and Oneida Lake for both the pre- and post-invasion periods. All information on lakes Eem and Veluwe was retrieved from Noordhuis et al. (2016).

Lake Lukomskoe is the 5$^{th}$ largest lake in Belarus with a wide littoral zone covered predominantly with sandy or silty sand sediments and occasionally with silt, and a profundal zone (> 6 m) with silt (Karatayev 1983). In 1969, Lake Lukomskoe became a cooling reservoir of the Lukoml Thermal Power Station. Zebra mussels were first found in this lake in 1972; no *Dreissena* were found during benthic studies in 1968 and 1969 (Lyakhnovich et al. 1982). Pre-invasion data for Lake Lukomskoe are limited. Before zebra mussel invasion, the lake was eutrophic with summer Secchi depth of 1.8 m in 1955 and 2.0 m in 1965 (reviewed in Lyakhnovich et al. 1988). Phytoplankton summer biomass for the pre-invasion period was estimated as 10.9 $gm^{-3}$, macrophyte coverage as 5% of the lake surface area, zooplankton biomass at 1.7 $gm^{-3}$ and zoobenthos biomass (excluding molluscs) as 0.6 $gm^{-2}$ (review in Lyakhnovich et al. 1988). Following the invasion of *Dreissena* in late 1970s - early 1980s, the lake became meso-oligotrophic (Karatayev 1983, Lyakhnovich et al. 1988). However, after the establishment of fish hatchery in cages targeted at 1000 tons of fish production per year, 3000 tons of organic matter with large amount of phosphorous (about 1% of fish food) started entering the lake, causing water quality deterioration, eutrophication, and oxygen decline (Mitrakhovich et al. 2008).

To evaluate *Dreissena* population dynamics and their ecological impacts on Lake Lukomskoe, we analyzed data available for the following years: Secchi depth: 1972, 1973-1981, 1983-1985, 1987, 1989-1992, 1994, 2003-2005, 2008, phytoplankton biomass: 1972, 1974-1979, 1983, 1984, 1987, 1989-1992, 1994, 2004-2005, macrophyte coverage: 1973, 1975, 1976, 1978, 1989, 1992, 2006, zooplankton biomass: 1972, 1974-1984, 1989-1992, 1994, 2003-2005, biomass of benthic invertebrates excluding molluscs: 1972, 1974-1978, 1981-1983, 1985, 1987 1989-1992, 2003-2005, and *Dreissena* lake-wide biomass: 1972, 1974-1978, 1989, and 2005 (Karatayev 1983; Karatayev et al. 1997; Lyakhnovich et al. 1982, 1988, Mitrakhovich et al. 2008).

Oligo-mesotrophic Lake Naroch and mesotrophic Lake Myastro were colonized by zebra mussels in mid-1980s (Burlakova et al. 2006). These lakes are among the best-studied lakes in the former Soviet Union. Data on water chemistry and major aquatic communities (phytoplankton, zooplankton, and benthos) are available since 1978 (Winberg 1985, Ostapenya et al. 1993, 1994, 2012). Zebra mussels were found in 1984 in Lake Myastro (Burlakova 1998) and in 1989 in Lake Naroch (Ostapenya et al. 1993).

Oneida Lake at 207 km$^2$ is the largest lake within the borders of New York State. It is a shallow, polymictic, meso to eutrophic lake with some bays and shoal areas covered with macrophytes down to 6 m, although coverage varies with fetch and substrate (Fitzgerald et al. 2016). The bottom substrate is variable and consists of rocky, sandy and silty areas, with most of the bottom below 9 m in silt. Few mussels were found in this deeper area until the expansion of quagga mussels in 2008 – 2009 (Hetherington et al. 2019). The lake is an important fishing lake, with large populations and fisheries for walleye (*Sander vitreus*) and black bass (*Micropterus sp*.). The lake has been sampled for a variety of variables, for some as far back as 1956 (Rudstam et al. 2016). All data used in this paper are available online through the Knowledge Network for Biocomplexity (Rudstam 2020a, b, c, d, e) and these references also includes details of sampling and analysis methods. Analysis of the ecosystem effects of zebra mussels goes back to the early years of the invasion (Mellina et al. 1995, Horgan and Mills 1999, Idrisi et al. 2001, Mayer et al. 2002) and mussel effects are a central theme of several chapters of a 2016 book on Oneida Lake (Rudstam et al. 2016).

##### *Dreissena* sampling protocol

The most detailed *D. polymorpha* study in Lake Lukomskoe was conducted in the summer of 1978, when samples were collected from 14 transects regularly distributed throughout the lake running perpendicular to the shoreline (Karatayev 1983). At each transect samples were collected from 0.5, 1, 1.5, 2, 3, 4, 5, 6, and 8 m depth. From 0.5 to 6 m depth samples were collected by SCUBA divers using quadrat (0.25 m$^2$), and an Ekman grab (0.025 m$^2$ sampling area) was used at 8 m depth. In 1989 and 2005, *Dreissena* samples were collected from 7 of the 14 transects sampled in 1978 using quadrats and bottom grabs (Peterson grab on hard substrates and Ekman grab on soft unconsolidated sediments, both 0.025 $m^2$) (Karatayev 1983, Mitrakhovich et al. 2008). At each transect 8 – 10 stations were sampled at depths similar to those sampled in 1978. All pre-1978 *Dreissena* surveys were conducted using bottom grabs. We do not have the information on the amount of sampling stations for most of these years, however most likely they ranged between 20 and 60 stations per sampling event.

Detailed sampling protocol for *Dreissena* density and biomass assessment in the Narochanskie Lakes can be found in Burlakova et al. (2006). In brief, samples were collected in Lake Naroch from eight permanent transects in 1990, 1993, 1994, 1995, and 1997 (Fig. 1) and latter in 2005 (Mastitsky et al. 2006). In Lake Myastro samples were collected from 5 transects per lake in 1993 and 1995. For all years and all lakes, transects were initiated on the shore and ran perpendicular to the shore toward the center of the lake. For each transect, up to 10 replicate samples were collected at 0.5, 1, 1.5, 2, and 3 m depth and then at an interval of 1 or 2 m down to the maximum depth where zebra mussels were found. In 1990-1997 samples down to 2 m were collected using the quadrat method (0.25 $m^2$). In Lake Naroch in 1997 quadrat samples were collected down to 7 m depths with the aid of a surface supplied air diving system (Pioneer 230 X, Brownie's Third Lung). A similar sampling design was implemented in Narochanskie lakes in 2016-2018 when divers collected samples from 0.5 to 8 m depth using 0.25 $m^2$ quadrat. Deeper samples in all lakes were collected with an Ekman grab on soft sediments or a Peterson grab on hard sediments (both 0.025 $m^2$).

All quadrat and grab samples collected from all Belarusian lakes were washed through a 500 μm mesh, and within 48 h of sampling all zebra mussels larger than 1 mm maximum dimension were counted, opened with a scalpel to remove water from the mantle cavity, and the total sample was weighed to the nearest 0.01 g after being blotted dry on absorbent paper (wet weight, soft tissue plus shell).

Oneida Lake dreissenid sampling developed over time (details in Hetherington et al. 2019). Mussels were collected at 8 - 10 sites (1992-2002) and at 15-26 sites (2003-2013) by SCUBA divers. Triplicates were taken at each site (1992-2002) and at most sites (2003-2013). Since 2013, Ekman grabs (1-3 replicates per site) were used on soft and sandy substrates and the number of sites was increased. SCUBA divers sampled rocky substrates as in past years. Mussels were returned to the lab the same day and frozen until processing time. Mussels were then thawed, identified to species, counted, and measured. Shell-on wet weights were calculated based on the mid-point of the length group using the regression shell-on wet weight = exp(2.97*ln(L, mm)-8.99) derived for zebra mussels in 1993 (Rudstam 2020a). Whole lake abundance estimates are based on the proportion of the lake bottom in different substrate and depth regions (shallower than 9m – rock, sand and soft substrate, deeper than 9 m).

***Environmental variables***

Environmental variables analyzed in this paper as response variables (potentially affected by *Dreissena*) to evaluate the impacts of *Dreissena* spp. invasion included transparency (as estimated by Secchi depth, m), total phosphorus concentration (mg/L, not studied in Lake Lukomskoe), macrophyte coverage (% of the lake bottom), chlorophyll *a* (g/m$^3$, not studied in Lake Lukomskoe), and wet biomass of phytoplankton (g/m$^3$), zooplankton (g/m$^3$), and benthos excluding molluscs (g/m$^2$). In Oneida Lake benthos biomass was not determined and for the model density was used instead. Dry mass of zooplankton calculated for Oneida Lake was multiplied by 5 to obtain wet biomass.

<u>Lake Lukomskoe</u>

No information was available on the amount of sampling stations and sampling events per year for Lake Lukomskoe, especially for pre-1977 period. Starting in 1977, phytoplankton and zooplankton in the lake was sampled at two pelagic stations at 8.0 m depth from surface, 2, 4, 6, and 7.5 m depths, usually 3 – 10 times per growing season. Phytoplankton was sampled using 1L Ruthner bathometer. Immediately after collection 0.5 L samples were preserved with acid Lugol's solution and kept in amber glass bottles in dark for about 10 days. Then, the top layer of water was siphoned, and samples were concentrated to 100 mL.

Zooplankton was collected with 10 L Vovk sampler and filtered through 64 μm mesh. Samples were concentrated to ca. 150 mL and fixed with formalin. Zooplankon groups (cladocerans, copepods, and rotifers) were counted under a dissection scope and identified to the lowest practical level using compound microscope. Biomass was estimated by multiplying the density of each species by their individual standard wet weight (same weight was used for the whole time period).

Zoobenthos was sampled from 15 – 20 stations (occasionally up to 46 stations) 3 to 10 times per year and washed through a 500 μm mesh. All macroinvertebrates collected were fixed with 10% buffered formalin, identified to the lowest practical taxonomic level (usually species, genus or family), counted, blotted dry on absorbent paper and weighed to the nearest 0.1 mg (total wet mass).

<u>Narochanskie Lakes</u>

To study chlorophyll, phytoplankton, and zooplankton in Narochanskie lakes, samples were collected monthly since 1978 during the vegetation period (May–October) from standard monitoring stations: two pelagic stations in Lake Naroch, and one station in Lake Myastro. Samples were collected using two-liter Ruttner sampler from six depth layers (0.5, 3, 6, 8, 12, and 16 m) in Lake Naroch and from four depths (0.5, 4, 7 and 9 m) in Lake Myastro. Water samples from all the depths for each lake were mixed proportionally to the fraction of the depth layer in the total volume of the lake to obtain an integral sample representing the mean composition of lake water. Secchi disk transparency was measured with a 30 cm diameter white disk.

To determine chlorophyll concentration (without correction for the presence of pheopigments) lake water was filtered using “Nucleopore” nuclear membrane with a pore diameter 1.5 μm. Chlorophyll analysis was performed spectrophotometrically after extracting pigments in 90% acetone (SCOR-UNESCO, 1996; Kovalevskaya et al. 2020).

Phytoplankton samples (0.5 L) were fixed with Utermöhl’s solution (Mikheyeva 1989) and analyzed using Zeiss Axiolab microscope. A Fuchs-Rosenthal chamber with a volume of 3.2 $mm^3$ was used to count small phytoplankton; larger species were counted in a 1 mL chamber, while large colonial organisms (such as *Gloeotrichia echinulata*, *Volvox*) were counted using Bogorov chamber. The phytoplankton density was expressed in cell number (number of one-celled species, number of cells in filaments and colonies) per liter, and phytoplankton biomass was estimated using the biovolume of individual algal species assuming a specific density of 1 $g/cm^3$. Biovolumes were obtained by measuring cell sizes and comparing cell shapes with geometric figures (Hillebrand et al. 1999, Mikheyeva 1999).

For zooplankton samples, 10 L of water was filtered using 64 μm mesh size and fixed in 4% formalin. Individual zooplankton were identified to species or genera, counted and measured using Zeiss Axiolab and Zeiss Stemi 2000 microscopes. The body mass of Cladocera and Copepoda was determined using length/volume relationship assuming a specific density of 1 $g/cm^3$. Rotifer species body volume was calculated using geometric figures most comparable to each species’ body shape using equations in Balushkina and Winberg (1979). The specific density of zooplankton was set to 1 $g/cm^3$.

Zoobenthos samples were taken usually three times per year along a nearshore to offshore transect. For Lake Naroch sample depths along the transect ranged from 1 to 16 m and for Lake Myastro from 1 to 10 m. In lakes Naroch and Myastro, samples were collected at every two meters (a total of 9 stations were sampled in Lake Naroch, 6 stations in Lake Myastro). Samples were collected with Petersen grab on hard and Ekman-Burge sampler with the Borutsky modification on soft substrates (sampling areas of both grabs 0.025 $m^2$), washed through a 265 µm mesh, and fixed with 10% neutral buffered formalin. Macroinvertebrates were identified to the lowest possible taxonomic level (usually species, genus or family) and counted. The biomass of individuals or groups of zoobenthos was determined by weighing on a torsion balance after blotted dry on absorbent paper.

Oneida Lake

Oneida Lake limnology data has been collected since 1975 from four or five sites at weekly intervals from soon after ice-out through late fall. To make data comparable with other lakes (e.g., Naroch, Myastro), we used the average of weekly measures collected in May through October, while for comparison of zebra vs. quagga mussel impact in Oneida Lake for Secchi depth, chlorophyll and total phosphorous we used data collected in March through November. Integrated water samples were collected with a 1.9 cm inner diameter tygon tube (Nalgene) lowered from the surface to 0.5 m above the bottom, then closed at the top, retrieved, and emptied into 4L plastic bottles or into a plastic carbuoy. Within 2 to 4 h after collection, samples for chlorophyll *a* were filtered onto a Whatman 934-AH filter and frozen. Filters were extracted with 90% acetone and measured spectrophotometrically (Strickland and Parsons 1972). Secchi disk transparency was measured with a 20 cm diameter black and white disk.

Subsamples of integrated lake water were preserved in Lugol's solution (1975 to 1995) or glutaraldhyde (1996 to present). From 1975 to 1995, the phytoplankton samples were settled in Utermöhl settling tubes at 10 ml aliquots for bloom phase samples and 25 ml aliquots for clear water phase samples. The samples were left to settle over a 24 hr period, after which the settling slide was placed in a Wild inverted microscope for identification and enumeration. Identification was made at two different magnifications (400x and 100x) to the species level when possible; otherwise, phytoplankton cells were identified to the genus. Phytoplankton were identified to species or higher taxonomic level and abundances were reported in #/mL. Biovolumes for this time period was estimated from standard species specific values.

From 1996 onwards, phytoplankton samples from different stations were pooled prior to analyses and 8 to 14 weekly samples were counted by Phycotech Inc. No samples were processed from 1998 and 1999. Phycotech counted and measured 400 natural units including all algal cells that were alive at the time of sampling (cells with content) using magnifications of 250x, 500x and 1250x. The 500x magnification was the primary one used. The algae were mounted on slides and the amount of water mounted varied with the density of algal cells (0.1 to 100 mL/slide). Algal taxa were identified to species when possible and measured to provide sample specific biovolumes for the sample, by cell and by natural units (colonies, paired or other multiples of cells).

Zooplankton were collected from the same sites using a 153-um mesh nylon net (0.5 m diameter) towed vertically from approximately 0.5 m off the sediment surface to the water surface. The efficiency of the net was measured with flow meters from 1999 to present. If flow meter readings indicate a malfunction or human error (efficiencies below 50% and above 125%) and when no flow meter was used, we assumed an efficiency of 87.4% (average of the 1999 to 2010 sampling period 87.4% SD 9.5%, N=1655). Flow meters were calibrated each year. Samples were preserved in 8 % buffered sugar-formalin solution (1975-1996) or 70 % ethyl alcohol (1997-present). Crustacean zooplankton were identified to species, counted and measured. A minimum of 100 animals were counted and measured from each sample. Biomass for individual species was calculated using length-weight regressions based mainly on Bottrell et al. (1976) and summarized in Watkins et al. (2011). Rotifers were not counted in Oneida Lake.

Benthos samples were collected from three sites using a 152x152 mm square (0.023 $m^2$) Ekman grab sampler. Samples were passed through a 253 μm mesh screen, preserved in ethanol, and stained with rose bengal. Organisms were removed from samples without a hand lens or microscope, and categorized to various taxonomic levels, mostly family or higher level.
